# How do zebrafish respond to MK-801 and amphetamine? Relevance for assessing schizophrenia-relevant endophenotypes in alternative model organisms

**DOI:** 10.1101/2020.08.03.234567

**Authors:** Radharani Benvenutti, Matheus Gallas-Lopes, Adrieli Sachett, Matheus Marcon, Nathan Ryzewski Strogulski, Carlos Guilherme Rosa Reis, Rafael Chitolina, Angelo Piato, Ana Paula Herrmann

**Affiliations:** Department of Pharmacology, Institute of Basic Health Sciences, Federal University of Rio Grande do Sul (UFRGS), Av. Sarmento Leite, 500/305, Porto Alegre, RS, 90050-170, Brazil; Graduate Program in Neuroscience, Institute of Basic Health Sciences, Federal University of Rio Grande do Sul (UFRGS), Av. Sarmento Leite, 500/305, Porto Alegre, RS, 90050-170, Brazil; Department of Biochemistry, Institute of Basic Health Sciences, Federal University of Rio Grande do Sul (UFRGS), R. Ramiro Barcelos, 2600, Porto Alegre, RS, 90035-003, Brazil; Graduate Program in Pharmacology and Therapeutics, Institute of Basic Health Sciences, Federal University of Rio Grande do Sul (UFRGS), Av. Sarmento Leite, 500/305, Porto Alegre, RS, 90050-170, Brazil

**Keywords:** zebrafish, schizophrenia, dizocilpine maleate, dextroamphetamine, locomotion, social behavior

## Abstract

**Background and Purpose:** Schizophrenia pathophysiology has been associated with dopaminergic hyperactivity, loss of parvalbumin-positive GABAergic interneurons, NMDA receptor hypofunction, and redox dysregulation. Most behavioral assays and animal models to study this condition were developed in rodents, leaving room for species-specific biases that could be avoided by cross-species approaches. As MK-801 and amphetamine are largely used in mice and rats to mimic schizophrenia features, this study aimed to investigate the effects of these drugs in zebrafish.

**Experimental Approach:** Adult zebrafish were exposed to MK-801 (1, 5, and 10 μM) or amphetamine (0.625, 2.5, and 10 mg·L^-1^) and observed in paradigms of locomotor activity and social behavior. Oxidative parameters relevant to schizophrenia were quantified in brain tissue.

**Key Results:** MK-801 disrupted social interaction, an effect that resembles the negative symptoms of schizophrenia. It also altered locomotion in a context-dependent manner, with hyperactivity when fish were tested in the presence of social cues and hypoactivity when tested alone. On the other hand, exposure to amphetamine was devoid of effects on locomotion and social behavior, while increased lipid peroxidation in the brain.

**Conclusion and Implications:** Key outcomes induced by MK-801 in rodents were replicated in zebrafish, which suggests this species is suitable as an alternative model animal to study psychotic disorders. More studies are necessary to further develop preclinical paradigms with this species and ultimately optimize the screening of potential novel treatments.

## 1 INTRODUCTION

Although the pathophysiology of schizophrenia is not completely understood, it has already been linked to GABAergic, glutamatergic, and dopaminergic dysfunction (Grace & Gomes, 2019; McCutcheon et al., 2020). Clinical and preclinical studies have suggested that psychotic symptoms arise from dopaminergic hyperactivity in subcortical regions (Kesby et al., 2018; McCutcheon et al., 2018), which may be due to loss of fast-spiking parvalbumin-positive GABAergic interneurons, hypofunction of NMDA receptors (NMDAR) and oxidative stress (Cabungcal et al., 2013; Hardingham & Do, 2016; Konradi et al., 2011; Steullet et al., 2010). Animal models of schizophrenia overwhelmingly rely on rodents, which may lead to species-specific biases that could be avoided by cross-species approaches (Burrows & Hannan, 2016; Weber-Stadlbauer & Meyer, 2019). Recent publications have endorsed the use of zebrafish as an alternative model animal to study schizophrenia and screen for potential novel treatments (Bruni et al., 2016; Gawel et al., 2019; Leung & Mourrain, 2016). More studies, however, are necessary to determine whether zebrafish models can provide sufficient behavioral and biochemical resolution to assess complex traits associated with psychiatric disorders such as schizophrenia.

Zebrafish have a conserved neural architecture analogous to mammals, including cell types, neurotransmitters, and receptors. Although catecholaminergic neurons are not present in the zebrafish midbrain, populations of tyrosine hydroxylase-immunoreactive cells have been identified in the diencephalon and telencephalon (Parker et al., 2013; Rink & Guo, 2004; Rink & Wullimann, 2001), with the structure and function of such neurons similar between mammals and teleosts (Matsui, 2017). Furthermore, glutamatergic and GABAergic neurons exert an important regulatory role in the zebrafish brain (Parker et al., 2013). The presence of these neuronal systems and their organization indicate it is feasible to use zebrafish to investigate the impact of psychotropic drugs on behavior and neurochemistry.

Well-established animal models of schizophrenia use pharmacological tools to recapitulate aspects of the neurobiology and symptomatology of the condition. MK-801 (dizocilpine), for instance, is a non-competitive NMDAR antagonist, while D-amphetamine (AMPH) inhibits dopamine uptake across the synaptic and vesicular membranes, leading to dopamine efflux. These drugs thus mimic hypofunction of NMDAR and dopaminergic hyperactivity, respectively (Jones et al., 2011). Inhibition of NMDAR in humans and animals leads to behavioral effects that resemble the full spectrum of positive, negative and cognitive symptoms of schizophrenia (Coyle et al., 2012; Hardingham & Do, 2016; Krystal et al., 1994; Morris et al., 1986). In mice and rats, MK-801 causes hyperlocomotion, stereotypic behavior, decreased social interaction, sensorimotor gating deficits, and cognitive impairment (Jones et al., 2011). In zebrafish, a few studies have shown that MK-801 exposure causes hyperlocomotion (Franscescon et al., 2020; Menezes et al., 2015; Tran et al., 2016), social deficits (Zimmermann et al., 2016), and cognitive impairment (Cognato et al., 2012; Franscescon et al., 2020; Gaspary et al., 2018). The behavioral effects of AMPH administration, on the other hand, only recapitulate the positive symptoms of schizophrenia, which include delusions, hallucinations and agitation in humans (Krystal et al., 2005), and stereotypic behavior and hyperlocomotion in rodents (Featherstone et al., 2007; Tenn et al., 2005). Although AMPH exposure in zebrafish is known to be anxiogenic (Kyzar et al., 2013), it has not been extensively investigated in the context of psychosis in this species.

Although MK-801 and AMPH are widely used in rodent models, their effects on behavioral and neurochemical parameters in zebrafish need to be further investigated. This is an important step to develop high-throughput drug screening platforms that may facilitate the discovery of novel antipsychotic agents. Thus, this study aimed to investigate the effects of acute exposure to MK-801 and AMPH in adult zebrafish by analyzing locomotor activity, stereotypy-related behaviors, social behavior, and neurochemical parameters of oxidative status as translatable markers.

## 2 METHODS

All procedures were approved by the institutional animal welfare and ethical review committee at the Federal University of Rio Grande do Sul (approval #35525/2019). The animal experiments are reported in compliance with the ARRIVE guidelines 2.0 (Sert et al., 2020).

### 2.1 Animals

Experiments were performed using 288 male and female (50:50 ratio) short-fin wild-type zebrafish, 6 months old, and weighing 400 to 500 mg. Adult animals were obtained from a local commercial supplier (Delphis, RS, Brazil) and maintained in our animal facility (Altamar, SP, Brazil) in a light/dark cycle of 14/10 hours for at least 15 days before tests. Fish were kept in 16-L (40 × 20 × 24 cm) unenriched glass tanks with non-chlorinated water at a maximum density of two animals per liter. Tank water satisfied the controlled conditions required for the species (26 ± 2 °C; pH 7.0 ± 0.3; dissolved oxygen at 7.0 ± 0.4 mg·L^-1^; total ammonia at <0.01 mg·L^-1^; total hardness at 5.8 mg·L^-1^; alkalinity at 22 mg·L^-1^ CaCO_3_; and conductivity of 1500–1600 μS/cm) and was constantly filtered by mechanical, biological and chemical filtration systems. Food was provided twice a day (commercial flake food (Poytara®, Brazil) plus the brine shrimp *Artemia salina*). After the tests, animals were euthanized by hypothermic shock according to the AVMA Guidelines for the Euthanasia of Animals (Leary et al., 2020). Briefly, animals were exposed to chilled water at a temperature between 2 and 4 °C for at least 2 minutes after loss of orientation and cessation of opercular movements, followed by decapitation as a second step to ensure death.

### 2.2 Materials

(+)-MK-801 hydrogen maleate (CAS Number: 77086-22-7) and D-amphetamine hemisulfate salt (CAS Number: 51-63-8) were purchased from Sigma-Aldrich (St. Louis, MO, USA). All drugs were dissolved in water at the same conditions as the home tanks. Solutions were freshly prepared immediately before tests and were renovated half-way through the tests. Reagents used for biochemical assays were obtained from Sigma-Aldrich (St. Louis, MO, USA) and included 5,5′-dithiobis(2-nitrobenzoic acid) (CAS Number 69-78-3), thiobarbituric acid (CAS Number: 504-17-6) and trichloroacetic acid (CAS Number: 76-03-9). Absolute ethanol (CAS Number: 64-17-5) was obtained from Merck KGaA (Darmstadt, Germany).

### 2.3 Experimental design

The animals were exposed to a concentration curve of MK-801 or AMPH for all tests. For MK-801 experiments, animals were randomly allocated to the following experimental groups: control (H_2_O); 1 μM MK-801; 5 μM MK-801, or 10 μM MK-801 (n=12). These concentrations correspond to 0.337, 1.687, and 3.373 mg·L-1, respectively. For AMPH experiments, animals were randomly allocated as follows: control (H_2_O); 0.625 mg·L-1 AMPH; 2.5 mg·L-1 AMPH or 10 mg·L-1 AMPH (n=12). These concentrations correspond to 3.392, 13.568, and 54.274 μM, respectively. We opted to express drug concentrations in the units routinely used in the literature for these drugs. The exposure time and concentration curves were based in the literature (Kyzar et al., 2013; Tran et al., 2016; Zimmermann et al., 2016) and pilot studies from our group. The animals were allocated to the experimental groups following block randomization procedures to counterbalance the sex, the two different home tanks, and the test arenas between the groups. Different sets of animals were used for each experiment. Animal behavior was video recorded and analyzed with the ANY-Maze tracking software (Stoelting Co., Wood Dale, IL, USA) by researchers blinded to the experimental groups. All tests were performed between 08:00 and 12:00 a.m. The sex of the animals was confirmed after euthanasia by dissecting and analyzing the gonads. For all experiments, no sex effects were observed, so data were pooled together.

### 2.4 Locomotor activity test

The experimental design of the locomotor activity test is depicted in figure 1A. To assess the effect of drug exposure across time on locomotor behavior, animals were individually and sequentially placed in (1) a beaker with 200 mL of water for 20 min, (2) the test aquarium for 30 min to analyze basal locomotor activity, (3) a beaker with 200 mL of water or drug solutions (MK-801 or AMPH) at different concentrations for 20 min, and (4) the test aquarium for 60 min. The test aquarium consisted of glass tanks (24 × 8 × 20 cm) filled with water at the optimal conditions at a level of 15 cm. The water in the tanks was changed between animals to avoid interference from drug traces or alarm substances released by previously tested fish. As the vertical position of the animal represents ethologically relevant information for assessing motor and anxiety-like behaviors (Levin et al., 2007), the test apparatus was virtually divided into three equal horizontal zones (bottom, middle and top) for the front view video analyses (Marcon et al., 2018). The following locomotor and exploratory parameters were quantified across bins of 5 min: total distance traveled, and time spent in the upper zone.

**Figure 1.**
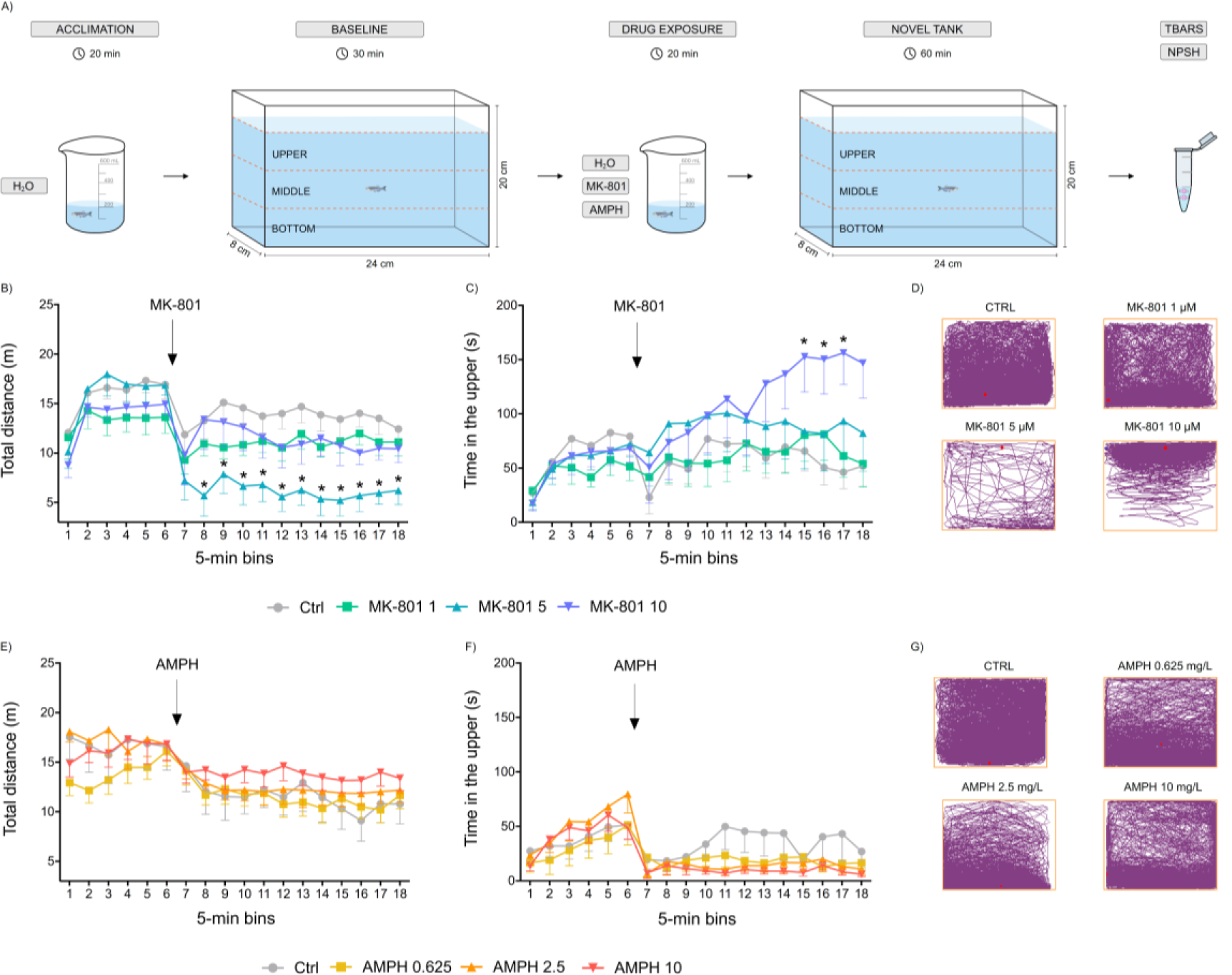
Effects of exposure to MK-801 and AMPH on locomotor activity across time. (A) Experimental design, (B, E) total distance traveled, (C, F) time spent in the upper zone, (D, G) representative track plots of one animal from each treatment group for 90 min. Data are expressed as mean ± standard error of the mean (S.E.M.). n=10-12. Repeated-measures ANOVA followed by Tukey’s post hoc test. *p<0.05 vs. control. AMPH (amphetamine); MK-801 (dizocilpine).

### 2.5 Open tank test

The experimental design of the open tank test is depicted in figure 2A. The apparatus used for the open tank test (OTT) consisted of a white circular arena (24 cm in diameter and 8 cm in height, 2 cm water level). The apparatus format and the acquisition of videos from the top view allows the evaluation of locomotor parameters related to stereotypic behavior (circular movements and absolute turn angle). For this test, the animals were individually exposed to water or different concentrations of the drugs in beakers filled with 200 mL of solution for 20 min. Following drug exposure, animals were placed in the center of the open tank arena and recorded for 10 min. The water was changed between every animal. For video analyses, the apparatus was virtually divided into two zones: the center zone of 12 cm in diameter and the periphery. As this experimental setup allows the acquisition of a great number of parameters, we performed a Principal Component Analysis (PCA) to investigate this dataset and selected for comparison only the parameters that contributed the most to total variation amongst animals (Fig. 6). The following parameters are thus displayed: total distance traveled, absolute turn angle, immobility time, number of clockwise rotations, and the time spent in the center zone.

**Figure 2.**
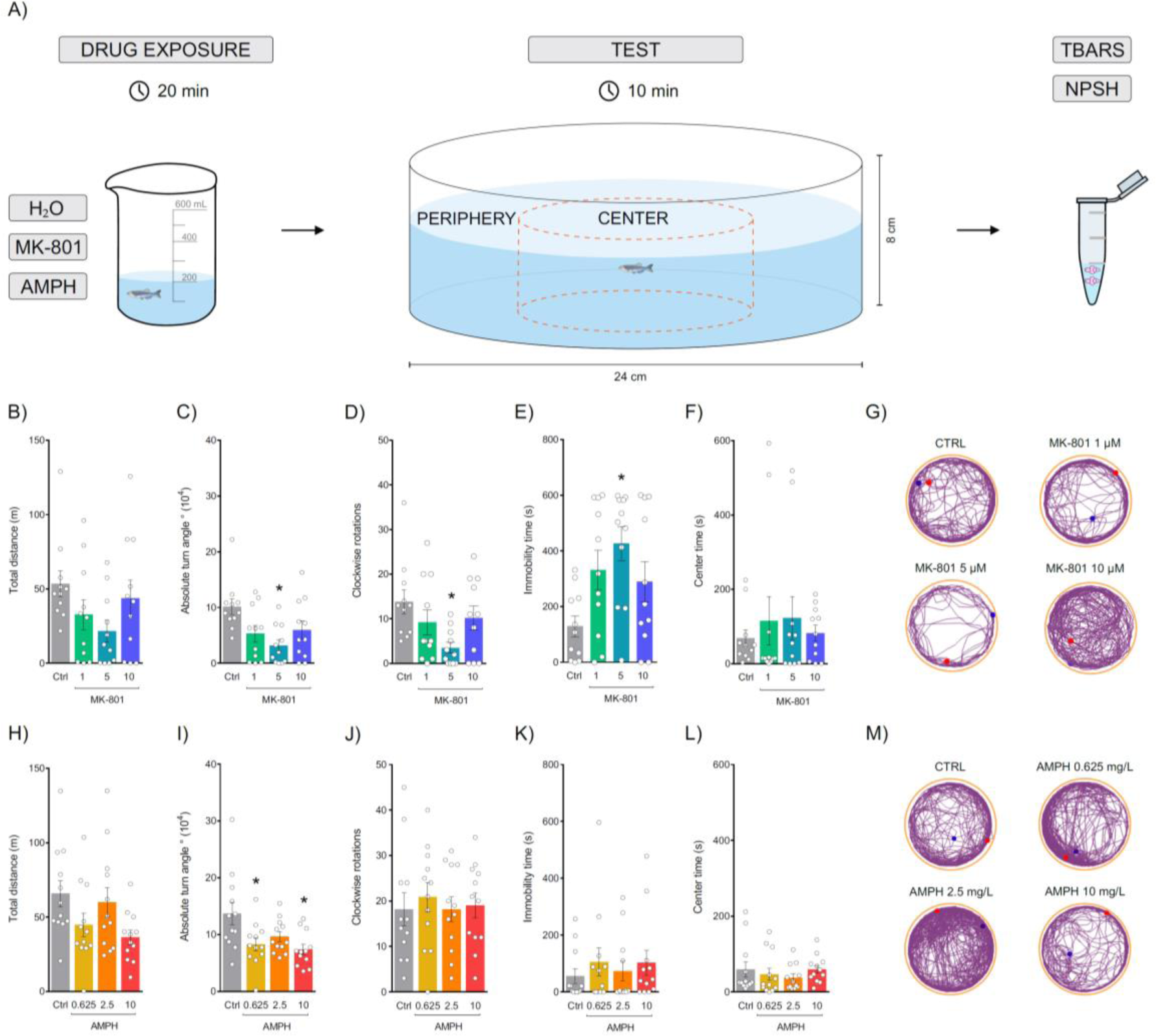
Effects of exposure to MK-801 and AMPH in the open tank test (OTT). (A) Experimental design, (B, H) total distance traveled, (C, I) absolute turn angle, (D, J) clockwise rotations, (E, K) immobility time, (F, L) time spent in the center zone, (G, M) representative track plots of the behavior of one animal from each treatment group during 10 min. Data are expressed as mean ± standard error of the mean (S.E.M.). n=11-12. One-way ANOVA followed by Tukey’s post hoc test. *p<0.05 vs. control. AMPH (amphetamine); MK-801 (dizocilpine).

**Figure 3.**
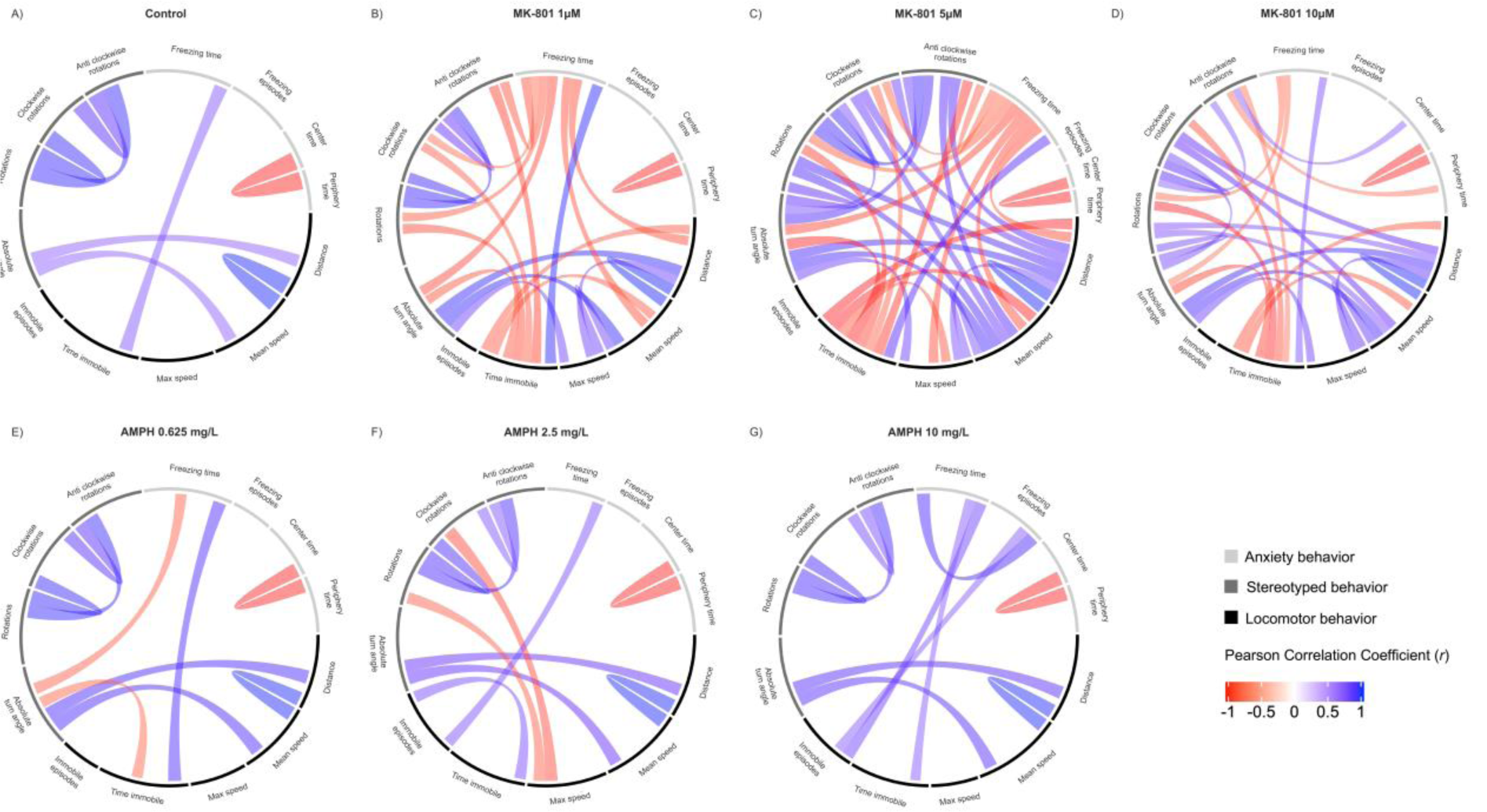
Mathematical analysis of the effects of MK-801 and AMPH on behavior during the open tank test (OTT). (A-G) Chord diagrams obtained from Pearson correlation analysis between variables, links indicate strong positive (blue color) and negative (red color) significant correlations, with “r” values above 0.7 or below -0.7, respectively. Annotation track colors indicate variables with related behavioral outcomes, classified as locomotor, stereotypy, and anxious behaviors, in black, gray, and light gray colors, respectively.

**Figure 4.**
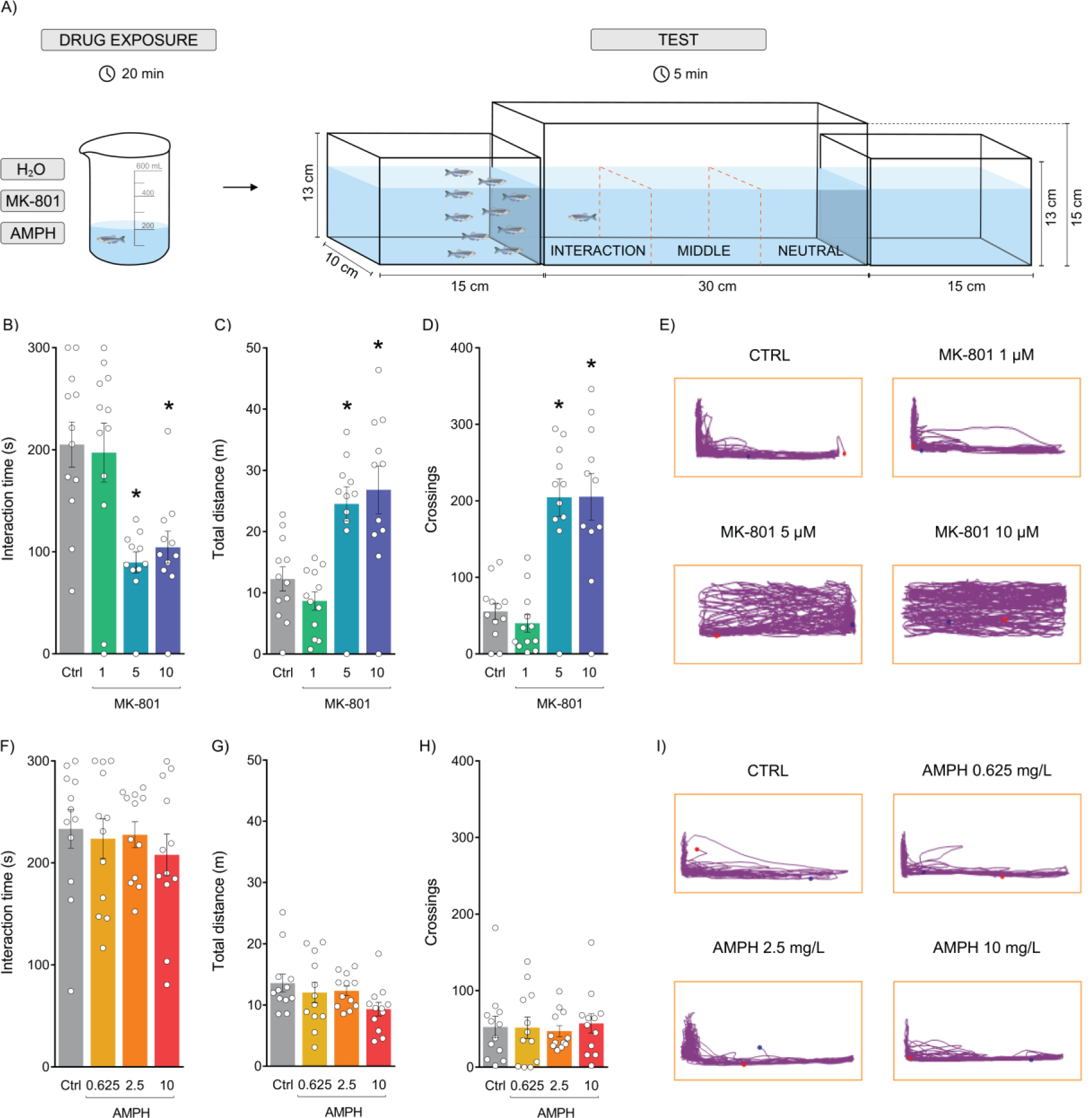
Effects of exposure to MK-801 and AMPH in the social interaction test. (A) Experimental design, (B, F) time spent in the interaction zone, (C, G) total distance traveled, (D, H) immobility time, (E, I) representative track plots of the behavior of one animal from each treatment group during 5 min. Data are expressed as mean ± standard error of the mean (S.E.M.). n =11-12. One-way ANOVA followed by Tukey’s post hoc test. *p<0.05 vs. control. AMPH (amphetamine); MK-801 (dizocilpine).

**Figure 5.**
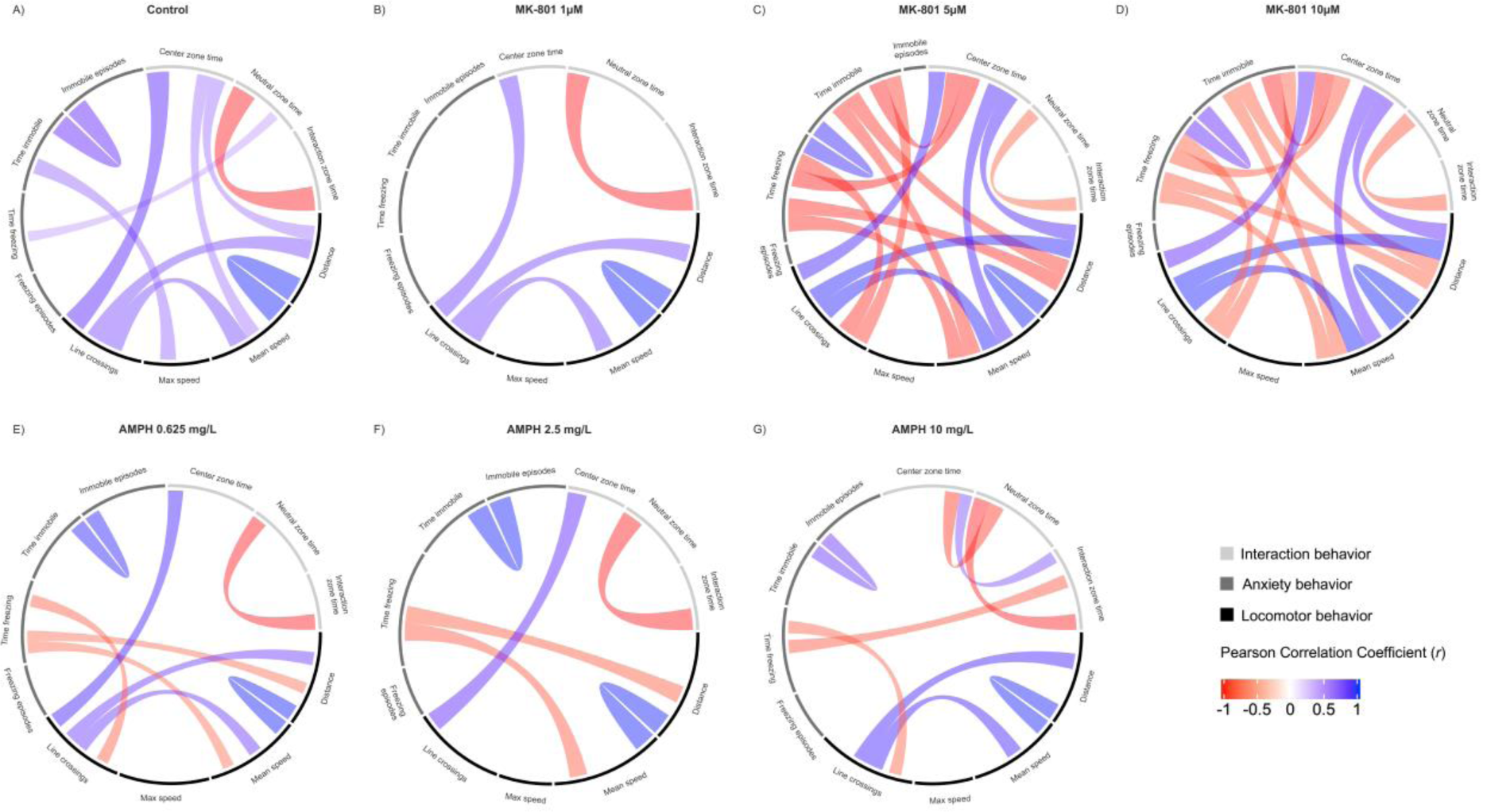
Mathematical analysis of the effects of MK-801 and AMPH on behavior during social interaction test. (A-G) Chord diagrams obtained from Pearson correlation analysis between variables, links indicate strong positive (blue color) and negative (red color) significant correlations, with “r” values above 0.7 or below -0.7, respectively. Annotation track colors indicate variables with related behavioral outcomes, classified as locomotor, anxiety, and social behaviors, in black, gray, and light gray colors, respectively.

**Figure 6.**
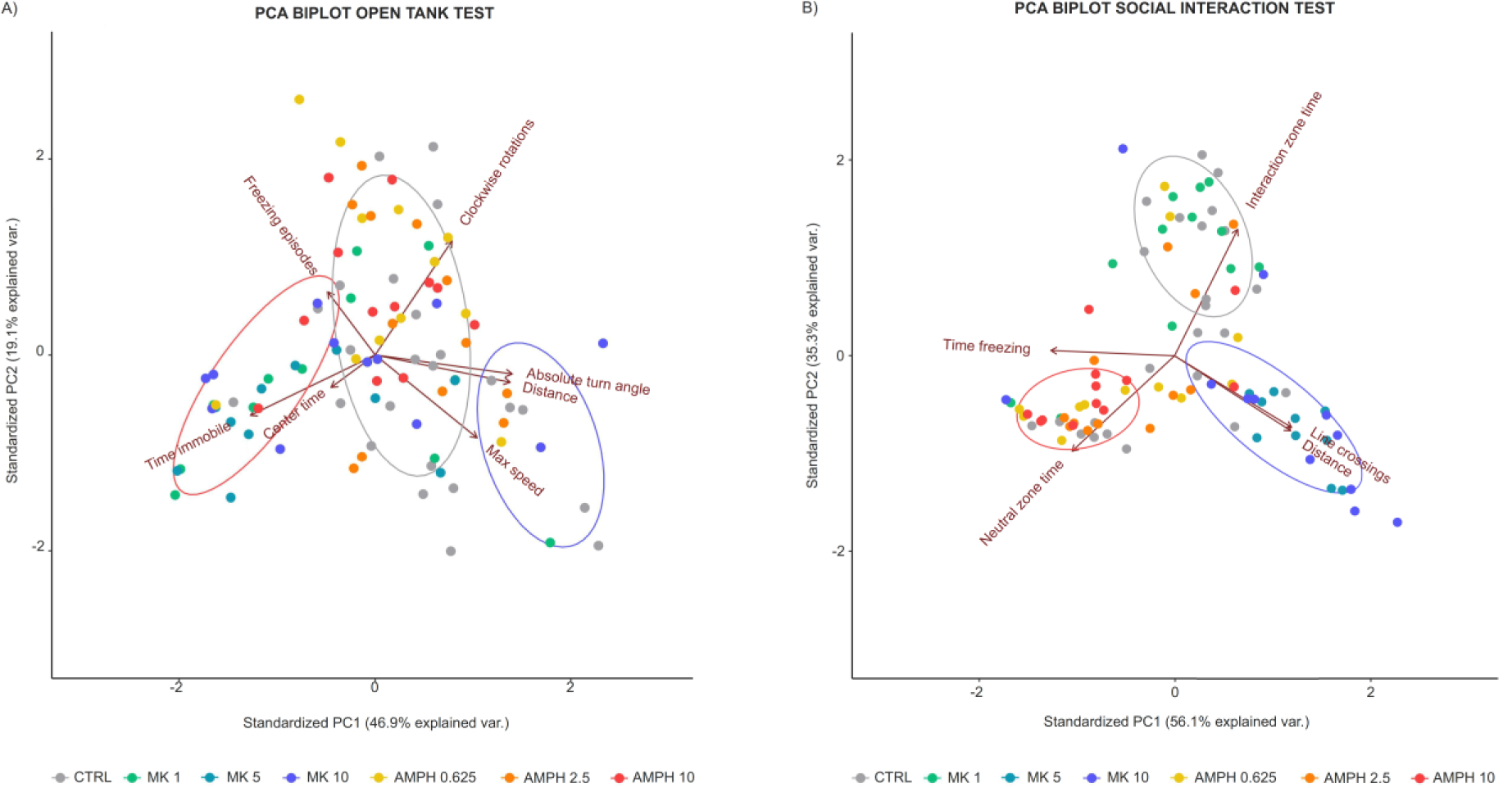
Mathematical analysis of MK-801 and AMPH effects on behavior during the open tank and social interaction tests. (A, B) Principal component analysis biplot displays arrows corresponding to observed variable eigenvectors relative to PC1 and PC2, grouping of animals in biplot is indicated in the legend. Ellipses indicate behavioral phenotype clustering model estimated through K-Means clustering analysis.

### 2.6 Social interaction test

The experimental design of the social interaction test is represented in figure 4A. In the social interaction test, animals were individually exposed to 200 mL of water or drug solutions in beakers for 20 min. After exposure, animals were placed for 7 min in a tank (30 × 10 x 15 cm) flanked by two identical tanks (15 × 10 x 13 cm) either empty (neutral stimulus) or containing 10 zebrafish (social stimulus). All three tanks were filled with water in standard conditions at a level of 10 cm. The position of the social stimulus (right or left) was counterbalanced throughout tests. The water in the experimental tanks was changed between every animal. To assess social behavior, the test apparatus was virtually divided into three vertical zones (interaction, middle and neutral). Videos were recorded from the front view. Animals were habituated to the apparatus for 2 min and then analyzed for 5 min. As for the open tank test, a large number of parameters may be obtained from this setup, so we also performed a PCA to investigate this dataset and selected for comparison only the parameters that contributed the most to total variation amongst animals (Fig. 6). The following parameters were quantified: total distance traveled, time spent in the interaction zone (as a proxy for social interaction time), and number of line crossings.

### 2.7 Oxidative stress parameters

The brain of animals submitted to the locomotor activity test and open tank test were collected right after the euthanasia to assess oxidative stress parameters by thiobarbituric acid reactive substances (TBARS) and non-protein thiol (NPSH) levels. A scalpel was used to remove the cranium of the fish and to collect brain tissue. For each independent sample, two brains were pooled (n=6) and homogenized in 300 µL of phosphate-buffered saline (PBS, pH 7.4, Sigma-Aldrich). Tissue preparation and biochemical analysis followed protocols described in (Sachett et al., 2018).

Firstly, the homogenates were centrifuged at 2400 g for 10 min at 4 °C, and the supernatants were collected and kept in microtubes on ice until the assays were performed. Lipid peroxidation was evaluated by quantifying the production of TBARS. 50 µg of proteins from the sample were mixed with thiobarbituric acid 0.5% and trichloroacetic acid 2% (150 µL). The mixture was heated at 100 °C for 30 min. The absorbance of the samples was determined at 532 nm in a microplate reader. 1,1,3,3-Tetraethoxypropane 2 nmol/mL was used as the standard. The content of NPSH in the samples was determined by mixing equal volumes of the brain tissue preparation and 6% trichloroacetic acid, centrifuging the mix (2400 g, 10 min at 4 °C), and determining the absorbance of the supernatants at 432 nm.

### 2.8 Data and statistical analysis

Data were expressed as mean ± S.E.M. For all comparisons, the significance level was set at p<0.05. Data were analyzed using IBM SPSS Statistics version 27 for Windows and the graphs were plotted using GraphPad Prism version 6.0 for Windows.

The sample size was calculated as n=12 for the primary outcome (total distance travelled) using MiniTab^®^ software. We used the following parameters: number of experimental groups (4), alpha (5%), power (80%), minimum difference between the means (50%) and standard deviation estimate based on the literature (40%).

The normality and homogeneity of variance confirmation were analyzed using D’Agostino-Person and Levene tests, respectively. The locomotor activity across time was analyzed using repeated-measures ANOVA to identify the main effects of treatment and time and their interaction, followed by Tukey post hoc test when appropriate. Data from the open tank test, social interaction test, and biochemical assays were analyzed using one-way ANOVA followed by Tukey post hoc test when appropriate. The outliers were defined using the ROUT statistical test and were removed from the analyses. This resulted in 2 outliers (1 from 5 μM MK-801 and 1 from 10 μM MK-801) removed from the social interaction test and 4 outliers (1 control animal and 1 from each MK-801 groups) removed from the open tank test. Also, 4 animals were excluded from locomotor activity test analyses due to technical problems in recording the videos (2 animals from the control group and 2 from 0.625 mg·L^-1^ AMPH) and 3 samples (1 sample from the control group and 2 samples from 0.625 mg·L^-1^ AMPH) were removed from oxidative stress analyses due to technical problems.

We performed Principal Component Analysis (PCA) to integrate the multiple variable outputs obtained during the open tank and social interaction tests. The new variables generated (principal components: PC’s) are a paired weighed combination of the variables contained in the original data that individually explain the largest possible variation amongst the samples. The PC’s were named in a decrescent manner, so that PC1 explains most variability, followed by the other PC’s in an ordered fashion of reduced variability explanation. Based on the composition of the PC’s and the individual contribution of each composing variable, it is possible to identify the most relevant and irrelevant variables to observe group differences. We also performed a K-Means clustering analysis (MacQueen, 1967), to quantitively assess the behavioral patterns observed in the Principal Component Analysis. Sampling adequacy was assessed before PCA analysis through Bartlett’s sphericity and Kaiser-Meyer-Olkin factor adequacy tests. Data frames presented p<0.05 for Bartlett’s sphericity test, and KMO of 0.62 and 0.66, respectively, indicating that both were adequate for PCA analysis. We estimated correlations using Person’s correlation test. We report only strong correlations (r^2^>0.5) and p<0.05. The statistical procedures were performed using base R-3.5.1 for MacOS (R Core Team, 2018), and R packages: “corrplot” (Wei & Simko, 2017), “psych” (Revelle, 2018), “circlize”(Gu et al., 2014), “ggplot2” (Wickham, 2016), “ggfortify” (Tang et al., 2016) and “devtools” (Wickham et al., 2019).

## 3 RESULTS

### 3.1 Effects of MK-801 and AMPH exposure in the locomotor activity test

In this experiment, we analyzed the effects of exposure to MK-801 (1, 5, and 10 µM) and AMPH (0.625, 2.5, and 10 mg·L^-1^) for 60 min, which allows a more detailed analysis of the locomotor activity and exploratory behavior of animals across time. As expected, no statistical differences were observed for baseline locomotor activity (Fig. 1B and 1E). However, after drug exposure, we observed that 5 µM MK-801 decreased the total distance moved (Fig. 1B) while 10 µM MK-801 increased the time spent in the upper zone (Fig. 1C). No statistical differences were observed after AMPH exposure in the parameters analyzed (Fig. 1E and 1F). Table 1 summarizes the repeated-measures ANOVA analyses.

**Table 1.** Summary of the repeated-measures ANOVAs for the locomotor activity test.

| Test | Dependent variable | Effects | F-value | DF | P-value | Effect size |
| --- | --- | --- | --- | --- | --- | --- |
| MK-801 locomotor activity | Total distance | Interaction | 2.63 | 51,748 | **<0.05** | 0.152 |
|  |  | Time | 14.88 | 17,748 | **<0.05** | 0.253 |
|  |  | MK-801 | 2.89 | 3,44 | **<0.05** | 0.165 |
|  | Upper time | Interaction | 1.87 | 51,748 | **<0.05** | 0.113 |
|  |  | Time | 6.62 | 17,748 | **<0.05** | 0.131 |
|  |  | MK-801 | 0.81 | 3,44 | >0.05 | 0.052 |
| AMPH locomotor activity | Total distance | Interaction | 1.07 | 51,680 | >0.05 | 0.075 |
|  |  | Time | 14.4 | 17,680 | **<0.05** | 0.265 |
|  |  | AMPH | 0.57 | 3,40 | >0.05 | 0.042 |
|  | Upper time | Interaction | 1.38 | 51,680 | **<0.05** | 0.094 |
|  |  | Time | 10.01 | 17,680 | **<0.05** | 0.200 |
|  |  | AMPH | 0.77 | 3,40 | >0.05 | 0.055 |

Significant effects ( $p < 0.05$ ) are given in bold font. Effect sizes are expressed as partial $\eta^2$ . DF, degrees of freedom.

### 3.2 Effects of MK-801 and AMPH exposure in the open tank test

In this experiment, we analyzed the effects of exposure to MK-801 (1, 5, and 10 µM) and AMPH (0.625, 2.5, and 10 mg·L^-1^) in the OTT, which allows the analysis of stereotypy-related behaviors, such as circular swimming and absolute turn angle. Exposure to 5 µM MK-801 decreased the absolute turn angle (Fig. 2C) and clockwise rotations (Fig. 2D) and increased the time spent immobile (Fig. 2E). Exposure to 0.625 and 10 mg·L^-1^ AMPH decreased the absolute turn angle (Fig. 2I) but did not alter clockwise rotations (Fig. 2J) or time spent immobile (Fig. 2K). There was no statistical difference in experimental groups regarding total distance (Fig. 2B and 2H) and time spent in the center area (Fig. 2F and 2L). Pearson correlation analysis shows that MK-801 (Fig. 3B, 3C, and 3D) exposure elicits a negative correlation between anxiety-related behaviors, namely freezing time, and locomotor and stereotypy-related behaviors, not observed in control animals (Fig. 3A). Interestingly, stereotypy- and locomotor-related behaviors were positively correlated in animals exposed to 5 µM MK-801 (Fig. 3C). A negative correlation between locomotor and stereotypy behaviors was only observed in animals exposed to 2.5 mg·L^-1^ AMPH (Fig. 3F). PCA analysis shows that PC1, corresponding to 46.9% of the explained variation, is largely positively composed of locomotor and stereotypy behaviors, whilst negatively composed of anxiety behaviors (Fig. 6A). A K-means analysis shows that 3 statistical clusters of similar behaviors are observed and that a cluster containing most of the animals treated with 5 µM MK-801 is observed, which is comprised of intense anxiety-like behaviors and reduced locomotor and stereotypy behaviors (Fig. 6A). Table 2 summarizes the one-way ANOVA analyses.

**Table 2.** Summary of the one-way ANOVAs for the open tank, social interaction, and oxidative status tests.

| Test | Dependent variable | F-value | DF | P-value | Effect size |
| --- | --- | --- | --- | --- | --- |
| MK-801 open tank test | Total distance | 1.00 | 3,40 | >0.05 | 0.070 |
|  | Absolute turn angle | 4.41 | 3,40 | **<0.05** | 0.249 |
|  | Clockwise rotations | 3.14 | 3,40 | **<0.05** | 0.191 |
|  | Immobility time | 2.67 | 3,40 | **<0.05** | 0.20 |
|  | Center time | 9.84 | 3,40 | >0.05 | 0.69 |
| AMPH open tank test | Total distance | 2.78 | 3,44 | >0.05 | 0.160 |
|  | Absolute turn angle | 4.46 | 3,44 | **<0.05** | 0.239 |
|  | Clockwise rotations | 0.16 | 3,44 | >0.05 | 0.11 |
|  | Immobility time | 0.35 | 3,44 | >0.05 | 0.24 |
|  | Center time | 0.54 | 3,44 | >0.05 | 0.036 |
| MK-801 social interaction test | Interaction time | 8.29 | 3,42 | **<0.05** | 0.372 |
|  | Total distance | 11.55 | 3,42 | **<0.05** | 0.452 |
|  | Crossings | 19.77 | 3,42 | **<0.05** | 0.582 |
| AMPH social interaction test | Interaction time | 0.36 | 3,44 | >0.05 | 0.24 |
|  | Total distance | 1.91 | 3,44 | >0.05 | 0.15 |
|  | Crossings | 0.117 | 3,44 | >0.05 | 0.008 |
| MK-801 oxidative stress | TBARS 10 min | 0.8639 | 3,20 | >0.05 | 0.115 |
|  | TBARS 60 min | 5.293 | 3,20 | **<0.05** | 0.442 |
|  | NPSH 10 min | 1.337 | 3,20 | >0.05 | 0.167 |
|  | NPSH 60 min | 0.7747 | 3,20 | >0.05 | 0.104 |
| AMPH oxidative stress | TBARS 10 min | 2.11 | 3,20 | >0.05 | 0.241 |
|  | TBARS 60 min | 8.783 | 3,19 | **<0.05** | 0.581 |
|  | NPSH 10 min | 0.2233 | 3,18 | >0.05 | 0.036 |
|  | NPSH 60 min | 0.9362 | 3,20 | >0.05 | 0.155 |

Significant effects ( $p < 0.05$ ) are given in bold font. Effect sizes are expressed as partial $\eta^2$ . DF, degrees of freedom.

### 3.3 Effects of MK-801 and AMPH exposure in the social interaction test

To assess behavioral parameters related to the negative symptoms of schizophrenia we investigated the effects of MK-801 and AMPH on zebrafish social behavior. Figure 4 shows the effects of exposure to MK-801 (1, 5, and 10 µM) and AMPH (0.625, 2.5, and 10 mg·L^-1^) in the social interaction test. 5 and 10 µM MK-801 decreased the time spent in the interaction zone (Fig. 4B), increased the total distance traveled (Fig. 4C) and increased the number of line crossings (Fig. 4D), while 1 µM MK-801 did not alter social and locomotor behaviors (Fig. 4B, 4C, and 4D). None of the AMPH concentrations altered social interaction (Fig. 4F) or locomotor behavior (Fig. 4G and 4H). Pearson correlation analyses showed that MK-801 exposure at 5 (Fig. 5C) and 10 µM (Fig. 5D) lead to negative correlations between anxiety and locomotor behaviors. AMPH exposure (Fig. 5E, 5F, and 5G) abrogated positive correlations between social behavior and both locomotor and anxiety behaviors, also leading to negative correlations between anxiety and locomotor behaviors. Interestingly, only 10 mg·L^-1^ AMPH exposed animals presented a negative correlation between anxiety behavior (time freezing) and time spent in the interaction zone (Fig. 5G). Further, PCA analysis shows that PC1, corresponding to 56.1% of the variation observed, is positively composed of total distance, crossings and time spent in the interaction zone, whilst negatively composed of time spent in the neutral zone and freezing time (Fig. 6B). PC2, corresponding to 35.3% of the variation observed is positively composed of time spent in the interaction zone and freezing time, whilst negatively composed of distance, crossings and time spent in the neutral zone (Fig. 6B). K-means analysis shows 3 distinct clusters among animals that suggest social interaction analysis in zebrafish presents a resolution to distinct phenotypes following 5 µM and 10 µM MK-801 exposure, associated with a reduced social interaction and increased locomotion (Fig. 6B). Table 2 summarizes the one-way ANOVA analyses.

### 3.4 Effects of MK-801 and AMPH exposure on oxidative status

Figure 7 shows the effects of exposure to MK-801 (1, 5, and 10 µM) and AMPH (0.625, 2.5, and 10 mg·L^-1^) on TBARS and NPSH levels. MK-801 exposure did not alter lipid peroxidation (Fig. 7A and 7B) and non-protein thiol levels (Fig. 7C and 7D) at any of the concentrations tested. AMPH, on the other hand, induced lipid peroxidation after 60 minutes of exposure (Fig. 7F), but not after 10 minutes (Fig. 7E), while it did not alter NPSH levels (Fig. 7G and 7H). Table 2 summarizes the one-way ANOVA analyses.

**Figure 7.**
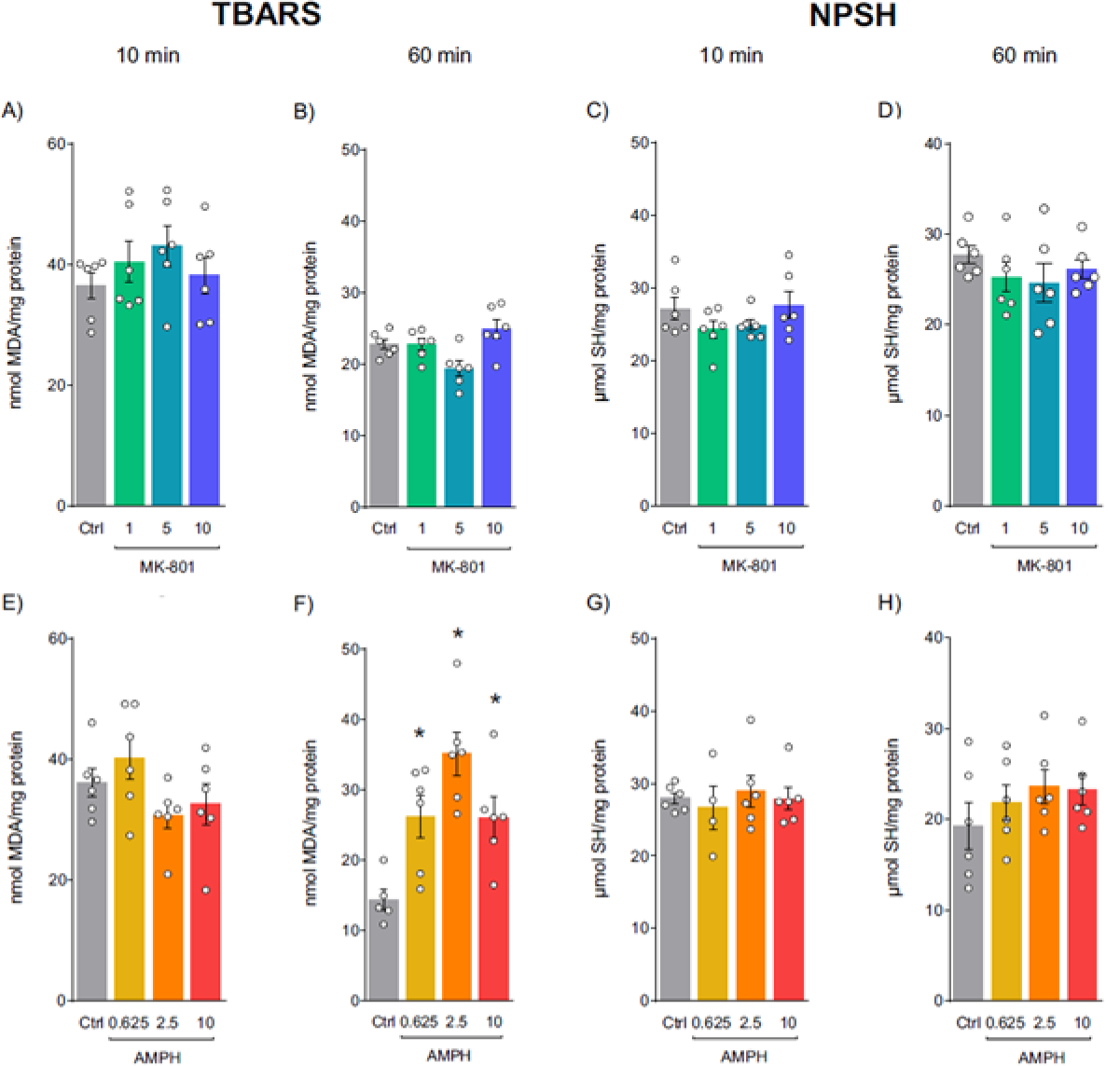
Effects of exposure to MK-801 and AMPH after 10 or 60 min on oxidative stress parameters (lipid peroxidation (TBARS) and non-protein thiol (NPSH) levels). (A, E) Lipid peroxidation levels 10 min after drug exposure, (B, F) lipid peroxidation levels 60 min after drug exposure, (C, G) NPSH levels 10 min after drug exposure, (D, H) NPSH levels 60 min after drug exposure. Data are expressed as mean ± standard error of the mean (S.E.M.). n=5-6. One-way ANOVA followed by Tukey’s post hoc test. *p<0.05 vs. control. AMPH (amphetamine); MK-801 (dizocilpine).

## 4 DISCUSSION

In the present study, we investigated the effects of exposure to MK-801 and AMPH on behavioral and biochemical outcomes in adult zebrafish. We demonstrate that MK-801 impaired social interaction, an endophenotype related to the negative symptoms of schizophrenia. Furthermore, we observed that MK-801 decreased the total distance traveled and increased the immobility time in locomotor assays, which differs from studies in rodents. Interestingly, MK-801 induced hyperlocomotion in the presence of social cues in the social interaction test. This suggests MK-801 elicits opposing effects that depend on the context, such as the presence of social stimulus. On the other hand, exposure to AMPH did not induce hyperlocomotion or social interaction deficit, but it was able to alter oxidative status, a relevant aspect related to schizophrenia pathophysiology.

Post-mortem brain studies and animal models of schizophrenia (Krystal et al., 1994; Mohn et al., 1999; Weickert et al., 2013) have suggested there is an aberrant function of hippocampal fast-spiking GABAergic parvalbumin-positive interneurons and a functional deficit in NMDAR expressed by these interneurons, which leads to altered excitation–inhibition balance in cortical and subcortical areas (Balu et al., 2013; Insel, 2010; Korotkova et al., 2010). MK-801 exposure simulates the NMDAR hypofunction, leading to disinhibition of excitatory hippocampal neurons and, consequently, disrupting the firing of dopaminergic neurons in the mesolimbic and mesocortical pathways, causing positive (hyperlocomotion and stereotypy-related behaviors) and negative (social interaction deficits) schizophrenia-like symptoms in rodents (Carlsson & Carlsson, 1989; Hardingham & Do, 2016). Exposure to AMPH increases dopamine release and is used as a pharmacological tool to model dopaminergic hyperactivity in the striatum (Featherstone et al., 2007; McCutcheon et al., 2020).

A behavioral assay widely evaluated in rodents is the distance traveled in an open field after intraperitoneal injection of MK-801 (Bygrave et al., 2016) or AMPH (Herrmann et al., 2014). Although the effects of these drugs on rodent locomotion are well known, the zebrafish literature is less straightforward. We thus performed in zebrafish a similar protocol to that used in rodents, in which locomotor activity is assessed across time. We also exposed the animals to an aquarium identical to the test aquarium before drug exposure to assess the basal locomotor activity, which in rodents declines and then stabilizes in the first 30 minutes. This decline in exploration was not observed for zebrafish in our study. We also surprisingly observed that 5 µM MK-801 decreased the total distance traveled, and 10 µM MK-801 increased time spent in the upper zone, which could indicate decreased anxiety. Contrary to our findings, some studies demonstrated that MK-801 exposure induced hyperlocomotion in adult zebrafish (Franscescon et al., 2020; Menezes et al., 2015), however such protocols lacked the baseline period and animals were directly placed in the test apparatus only after drug exposure. The differences in context novelty may underly the conflicting results, as suggested by a previous study by Tran et al. (2016).

In our experiments, AMPH did not induce hyperlocomotion in any behavioral test, which contrasts with the well-known stimulating effects observed in rodents. Differences in fish central regulatory motor circuits as compared to rodents and other mammals may explain our findings (Ryczko et al., 2017). An obvious difference, for example, is the fact that fish are swimming most of the time. Studies performed in lampreys (*Petromyzon marinus*) have shown that there are dopaminergic and glutamatergic neurons that project to the mesencephalic locomotor region (MLR), indicating a close interaction between these neurotransmitters in the generation of the locomotor command. Besides, blocking dopamine receptors in the MLR resulted in reduced swimming movements without interrupting the gradual locomotion control, while blocking glutamatergic receptors almost abolished locomotion, indicating that glutamatergic contribution is essential to obtain locomotion in a graded fashion, while the dopaminergic contribution provides additional modulation, but is not essential to evoke locomotion (Ryczko et al., 2017). We hypothesized that, because of the different central circuits of motor regulation of fish, the results with MK-801 exposure on zebrafish locomotion parameters are more robust than with AMPH exposure.

Stereotypy is defined as repetitive and unvarying behavior (Morrens et al., 2006). Drug-induced stereotypic behaviors are not well characterized in zebrafish, but some authors suggest they may be displayed as altered rotation movements (Michelotti et al., 2018). To date, only a few studies explored the effects of MK-801 on stereotypy-related behaviors in zebrafish, and there is no study with AMPH. Franscescon et al. (2020) report a decrease in the absolute turn angle after exposure to MK-801; however, the test aquarium in this study was filmed from the front and not from the top view, which hinders proper evaluation of circular movements in the horizontal plane of the fish. Another study found increased circular movements in zebrafish after exposure to a high concentration of MK-801 (100 µM) in the open tank test (Sison & Gerlai, 2011). In our study, 5 µM MK-801 decreased the absolute turn angle and clockwise rotations, and increased the time animals remained immobile, whereas 0.625 and 10 mg·L^-1^ AMPH only decreased the absolute turn angle. It is also possible that zebrafish display other phenotypes related to stereotypic behavior that are not detectable by automated software, such as repetitive movements of the face and mouth commonly observed in humans and rodents exposed to AMPH or NMDAR antagonists (Kelley et al., 1988; Ridley & Baker, 1982). We observed that in both MK-801 and AMPH exposures, stereotypy-related behaviors were positively correlated to locomotor-related parameters in the open tank test.

As social isolation is one of the main negative symptoms of schizophrenia, behavioral assays that model this aspect are critical for studying and developing treatments to this unmet clinical need. As zebrafish is a schooling animal with complex social behavior, such as hierarchical and breeding relationships (Dreosti et al., 2015), it is well-suited as a species for modeling social interaction. We observed that 5 and 10 µM MK-801 disrupted social interaction as measured by a decrease in the time zebrafish spent in the interaction zone. This result corroborates previous findings in other zebrafish studies (Seibt et al., 2011; Zimmermann et al., 2016) and in rodent models. This reinforces the notion that zebrafish is an adequate species to model an endophenotype related to the negative symptoms of schizophrenia. Interestingly, 5 and 10 µM MK-801 caused hyperlocomotion in the social interaction test, even though this drug was devoid of effects on the total distance traveled in the locomotor tests discussed earlier. This suggests that not only context novelty (Tran et al., 2016), but also the presence of social cues can differentially modulate the effects of MK-801. This idea, however, remains to be tested in experiments that isolate both variables. Although hyperlocomotion could be a confounder to social interaction, we observed that distance traveled did not correlate with time spent in the interaction zone, corroborating to the non-stochastic nature of this behavior. Regarding AMPH, it did not cause significant changes in zebrafish social behavior, replicating what is known for rodents. PCA and clustering analysis suggest that in fact, both open tank and social interaction tests have sufficient resolution to identify distinct behavioral phenotypes related to schizophrenia in zebrafish models, specially following MK-801 exposure.

Studies in rodent models report that increased dopamine release induced by AMPH promotes oxidative stress, probably due to increase monoamine metabolism (Dichtl et al., 2018; El-Tawil et al., 2011; Frey et al., 2006). All tested concentrations of AMPH increased TBARS, a well-established marker of lipid peroxidation, after 60 min, but not after 10 min of exposure, indicating a time-depend effect on the production of reactive species. This increase in TBARS levels, however, was not compensated by changes in NPSH levels, which indirectly represent reduced glutathione (GSH) levels and antioxidant activity. These are novel data since there are no studies evaluating the effects of AMPH on oxidative stress parameters in zebrafish. On the other hand, MK-801 did not cause any change in the oxidative status after 10 or 60 min of exposure. This is in line with studies in rodents that demonstrate that only repeated exposures to MK-801 produce an increase in reactive oxygen species (Wang et al., 2012).

With ongoing efforts in modulating schizophrenia-associated genes in zebrafish, straightforward behavioral readouts relevant to this condition need to be investigated. This is essential to advance the use of zebrafish in high-throughput drug screening assays. Our study corroborates the idea that schizophrenia endophenotypes may be modeled in lower organisms. In conclusion, MK-801 may be more useful than AMPH as a pharmacological tool to assess translatable behavioral markers relevant to schizophrenia in adult zebrafish.

## ACKNOWLEDGMENTS

This work was supported by Coordenação de Aperfeiçoamento de Pessoal de Nível Superior (CAPES) with fellowships to R.B., C.G.R.R., and R.C. We also thank the support from Conselho Nacional de Desenvolvimento Científico e Tecnológico (CNPq), and Pró-Reitoria de Pesquisa (PROPESQ) at Federal University of Rio Grande do Sul (UFRGS).

